# Dynein and dynactin move long-range but are delivered separately to the axon tip

**DOI:** 10.1101/2023.07.03.547521

**Authors:** Alexander D Fellows, Michaela Bruntraeger, Thomas Burgold, Andrew R Bassett, Andrew P Carter

## Abstract

Axonal transport is essential for neuronal survival. This is driven by microtubule motors including dynein, which transports cargo from the axon tip back to the cell body. This function requires its cofactor dynactin and regulators LIS1 and NDEL1. Due to difficulties imaging dynein at a single-molecule level, it is unclear how this motor and its regulators coordinate transport along the length of the axon. Here we use neuron-inducible human stem-celllines (NGN2-OPTi-OX) to endogenously tag dynein components and visualise them at a near-single molecule regime. In the retrograde direction, we find that dynein and dynactin can move the entire length of the axon (>500 *μ*m) in one go. Furthermore, LIS1 and NDEL1 also undergo longdistance movement, despite being mainly implicated with initiation of dynein transport. Intriguingly, in the anterograde direction, dynein/LIS1 move faster than dynactin/NDEL1 consistent with transport on different cargos. Therefore, neurons ensure efficient transport by holding dynein/dynactin on cargos over long distances, but keeping them separate until required.

## Introduction

The axon relies on a network of microtubule motors and associated proteins to maintain neuronal function. These factors transport cellular components such as RNAs, proteins, organelles and neurotrophic signals throughout the highly polarised neuronal environment^1^. Breakdown or impairment in these transport mechanisms is detrimental to the health of the cell with mutations and deficits linked to a range of neurological diseases^2^. The two main classes of motor proteins involved with transport in the axon are kinesin and dynein. Due to the organisation of axonal microtubules, kinesin motors drive cargos towards the distal tip (the anterograde direction), whereas a single dynein (cytoplasmic dynein-1, hereafter dynein) transports them back to the cell body (the retrograde direction)^1^. Dynein relies on its co-factor dynactin, cargo-specific activating adaptors and associated regulators such as LIS1 and NDEL1 to form a motile complex^3^. However basic questions about how dynein and its co-factors behave in the neuron, such as how far they move and whether they do so together, remain unanswered.

Dynein is ubiquitously expressed at high levels throughout the neuron^4^. Consequently, single molecules of motile dynein have proven difficult to visualise above the background of freely diffusing motor. Previous studies of neurons from a mouse expressing a GFP-tagged dynein intermediate chain used local photobleaching to observe movements of up to 15 *μ*m ^4,5^. In contrast, a recent study in HeLa cells, used highly inclined and laminated optical sheet (HILO) imaging to visualise single molecules of GFP-tagged dynein heavy chain. This suggested dynein has a short residence time on microtubules and only undergoes short-range (1-2 *μ*m) movements^6^, leading to the conclusion that long-range transport is achieved by a constant exchange of motile dynein complexes. These studies raise the question of how far dynein motors move in the axon. Can a single motor travel the whole distance (typically greater than 500 *μ*m) from the axon tip back to the cell body, or do cargos continuously replenish their pool of dyneins?

To elucidate how dynein drives long-range transport in neurons, we used human stem cell lines that can be differentiated into excitatory cortical neurons^7^. This enabled us to endogenously tag dynein and its associated proteins, avoiding any potential artefacts of overexpression^8^. We used HILO imaging ^6,9^ combined with SNAP-tag and HaloTag-linked fluorophores for improved photostability. This allowed live imaging of dynein molecules in human neurons at a near-single molecule regime. We discovered that both dynein and dynactin are highly processive along the entire length of the axon. Furthermore, we find that LIS1 and NDEL1, which are thought to play a role in initiation of dynein transport, also move long distances. Unexpectedly, when analysing the anterograde transport of dynein and dynactin we found that they display different average velocities, suggesting the majority of these proteins are transported separately towards the distal axon. Taken together, our study allows us to understand how the dynein machinery drives long-range transport of cargo in the axon.

## Results

### iNeurons as a model to study axonal transport

To study axonal transport, we used engineered human embryonic stem cell (hESC) or human induced pluripotent stem cell (hiPSC) lines. These contain a doxycycline-inducible, neurogenin 2 (NGN2) expression cassette in the adeno-associated-virus integration site 1 (AAVS1) safe harbour locus^7^. Upon treatment with doxycycline, stem cells undergo rapid, homogenous and highly reproducible differentiation into excitatory cortical neurons (hereafter referred to as iNeurons)^10,11,12^. In agreement with previous reports of NGN2 driven differentiation, cells begin to display clear neuronal morphology 7 days post induction (DPI) (Fig S1A)^7,11,13,14^. By 21-23 DPI cells have clearly-defined axons and dendrites as shown by immunofluorescence staining by SMI-31 and MAP-2 respectively (Fig S1B).

To assess axonal transport in iNeurons, cells were plated into microfluidic devices at 2 DPI and allowed to grow until 21-23 DPI^15^ (Fig 1 A,B). By this time point the axons have grown through the microfluidic grooves into the axonal compartment and are isolated from the dendrites. We then treated the axonal compartment with organelle-specific markers to label endosomes (Cholera toxin subunit B: CTB), lysosomes (lysotracker) and mitochondria (mitotracker) and characterised their movement (Fig 1C, Video 1-3). Endosomes displayed faster speeds than the other organelles, in agreement with previous observations in mouse/rat primary neuron cultures^16^ and another NGN2-induced neuronal model (i3 Neurons)^13^ (Fig 1D). Also in agreement with previous work, our endosomes and lysosomes moved primarily in the retrograde direction, whereas mitochondria displayed a high degree of bi-directional movement^13,17^ (Fig 1E). Given the similarities between our iNeurons and other previously reported models, we believe that they represent an excellent system to study the role of dynein in long-range transport.

**Figure 1:**
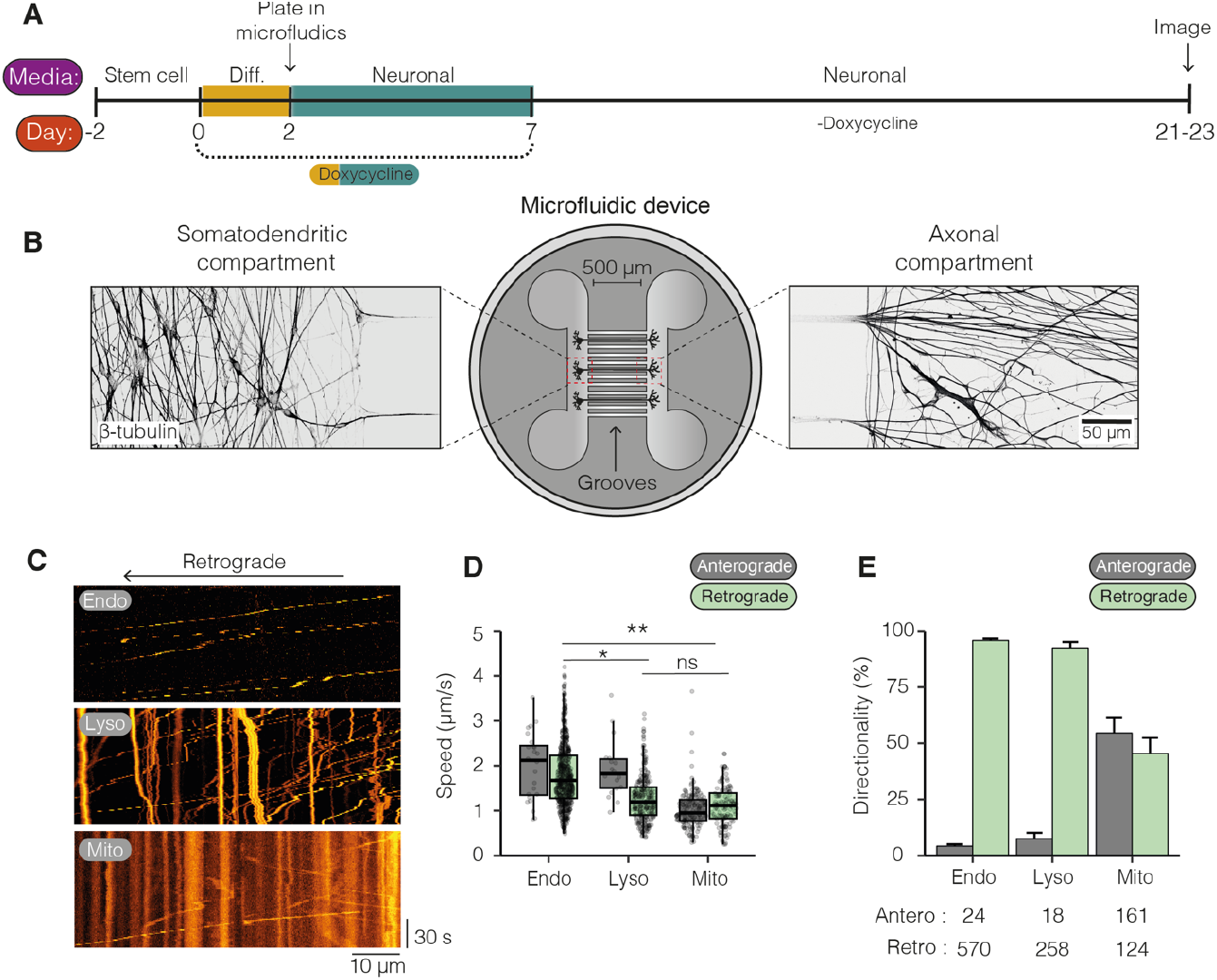
iNeurons as a model to study dynein-mediated transport. **A)** Schematic of differentiation of NGN2 hESCs/hiPSCs into iNeurons. hESC/hiPSCs were split and 300,000 cells were plated. Two days later differentiation media (Diff.) was added contain doxycycline. At 2 DPI, cells were split again and plated into microfluidics. At 7 DPI, doxycycline was removed from the media and cells were allowed to grow until 21-23 DPI.**B)** Example image of 21-23 DPI iNeurons in microfluidic device. Cells were fixed and stained with an antibody against *β*-tubulin. **C)** Kymographs of endosomes (Endo, CTB AlexaFluor 488), lysosomes (Lyso, Lysotracker Deep Red) and mitochondria (Mito, mitotracker Deep Red FM) in iNeurons at 21-23 DPI. **D)** The mean speed of endosomes, lysosomes and mitochondria in both anterograde (Grey) and retrograde (light green) directions (Retrograde: endo vs lyso *p=0.027, endo vs mito **p=0.0025, Kruskal-Wallis, Dunn post hoc test, N=3, stats fully reported in supplemental table 1). Boxplot shows median, first and third quartiles. Upper/lower whiskers extend to 1.5x the interquartile range. **E)** The directionality of endosomes, lysosomes and mitochondria movements in iNeurons at 21-23 DPI (Endosomes: 594 cargoes, 21 videos, N= 3; lysosomes: 276 cargoes, 11 videos, N=3; mitochondria: 285 cargoes, 22 videos, N=3). Error bars represent standard error of the mean (SEM).

### Visualising single dynein and dynactin molecules in iNeurons

To visualise motors with near-single molecule sensitivity we used CRISPR to endogenously tag the N-terminus of the dynein heavy chain with a HaloTag^18^ in our hESCs temp cell line (Halo-*DYNC1H1*, hESCs) and differentiated them into iNeurons. We labelled the HaloTag with Janelia fluor extra dyes (JFX554 or JFX650), which are brighter and more photostable than the GFP which was used previously to image dynein^4,19^. We first treated our Halo-*DYNC1H1* iNeurons with 1 nM JFX554 in the axonal compartment in order to label a subset of molecules for single molecule imaging as previously described for other systems^20^. We saw many distinct dynein spots in the axonal compartment, most of which were diffusing, often along microtubules (Fig S2A, Video 4). We observed rare instances of processive movement (Fig S2A. Video 4) consistent with previous studies suggesting only a subset of dyneins are actively involved in transport.

To ask whether our labelled dyneins are present as isolated molecules or as clusters we performed a photobleaching analysis (Fig 2A, B, C, Video 5,6). At first this proved difficult due the movement of the dynein spots in and out of the focal plane. Therefore we treated iNeurons with N-ethyl maleimide (NEM), a known inhibitor of motor proteins which traps them on microtubules^21,22^ (Fig 2 B,C). The immobilised dynein spots displayed between 1-7 clear photobleaching steps with 2 steps being the most common (Fig 2D, S2B, C). We repeated this analysis on iNeurons containing the endogenously tagged ARP11 subunit of dynactin (Halo-*ACTR10*, hiPSCs). We again saw between 1-7 photobleaching steps but now with 1 step begin the most frequent (Fig 2E). This difference in distribution of step sizes likely correlates with the fact there are two copies of the Halo-*DYNC1H1* in a dynein dimer, but only a single Halo-*ACTR10* per dynactin.

**Figure 2:**
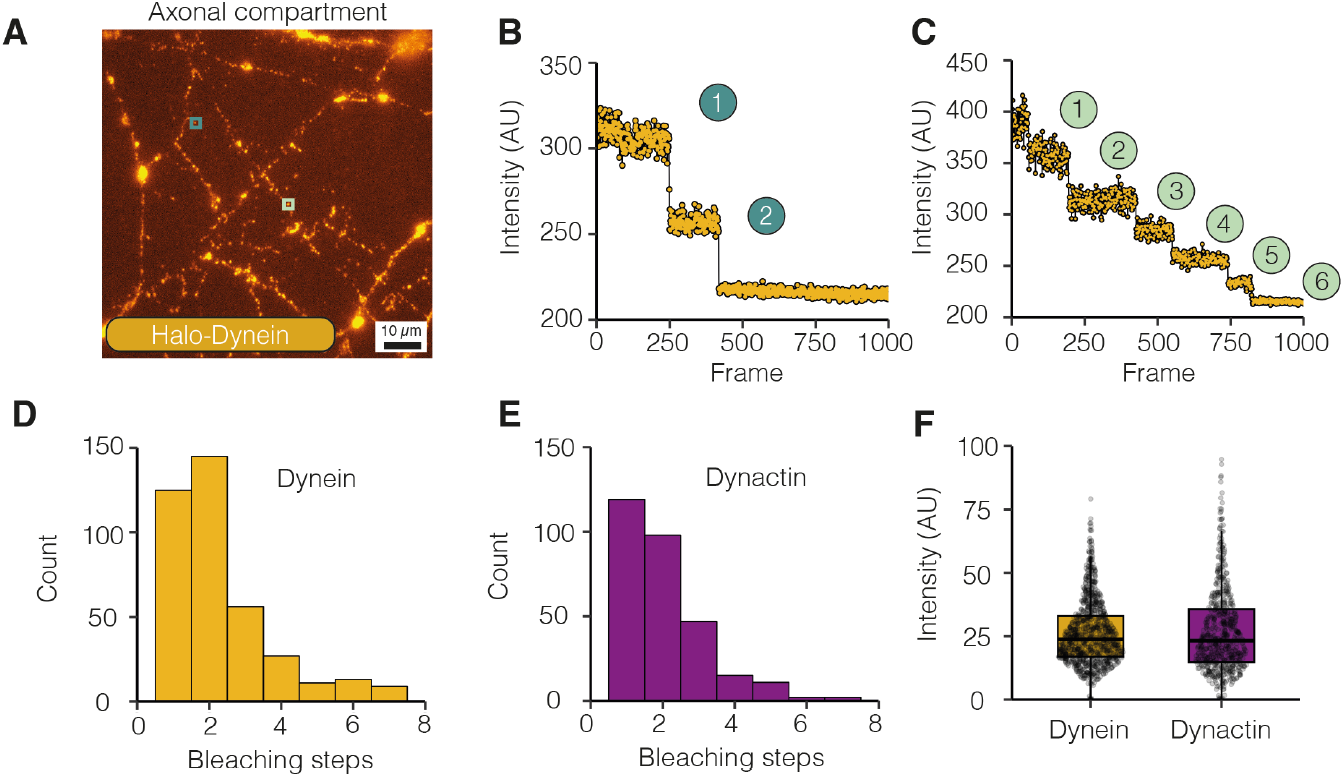
Visualising dynein in iNeurons. **A)** Image of 21-23 DPI Halo-*DYNC1H1* iNeuron axons stained with 1 nM JFX 554 and treated with 0.5 µM NEM. Teal and light green insets display spots before bleaching. **B)** The bleaching trace from teal inset in A. The spot displays two bleaching steps representing the presence of one dynein molecule. **C)** The bleaching trace from light green inset in A. The spot displays six bleaching steps representing the presence of three dynein molecule. **D)** The number of bleaching steps from dynein (Halo-*DYNC1H1*) spots (389 spots, 14 videos, N=3). **E)** The number of bleaching steps from dynactin (Halo-*ACTR10*) spots (297 spots, 10 videos, N=3). **F)** Graph shows the analysis of step size intensity from dynein and dynactin spots during bleaching (dynein: 912 steps, 14 videos, N=3; dynactin: 623 steps, 10 videos, N=3;). Boxplot shows median, first and third quartiles. Upper/lower whiskers extend to 1.5x the interquartile range.

Importantly, we also quantified the intensity of each step for both the dynein and dynactin photobleaching and found them to be very similar (dynein: 25.96 ± 4.38 AU, dynactin: 26.82 ± 6.08 AU, Fig 2F) suggesting they correspond to bleaching of individual fluorophores. Taken together our data imply that under these imaging conditions we are capable of detecting single molecules.

### Dynein moves long range

To directly address how far dyneins move we treated the axonal compartment of Halo-*DYNC1H1* iNeurons with 200 nM JFX 554/650. This concentration of dye labels dynein close to saturation ensuring as many dynein molecules were labelled as possible. After 20 min we collected movies in the microfluidic grooves near the somatodendritic compartment using the same imaging conditions as in previous experiments. Due to the fluidic isolation any observed fluorescent signal must travelled down the axon. When we imaged a control iNeuron line without any integrated HaloTag we saw no fluorescence at this time point (Fig S3A). In contrast when imaging Halo-*DYNC1H1* iNeurons we saw multiple highly processive spots moving predominantly in the retrograde direction (Fig 3A, Fig S3A, Video 7). The speed of these dynein spots ranged from 0.3-5.0 µm/s with an average of 1.76 ± 0.12 µm/s (Fig 3B), which agrees with speeds of retrograde organelles both in these neurons (Fig 1D) and in the literature^13^.

**Figure 2:**
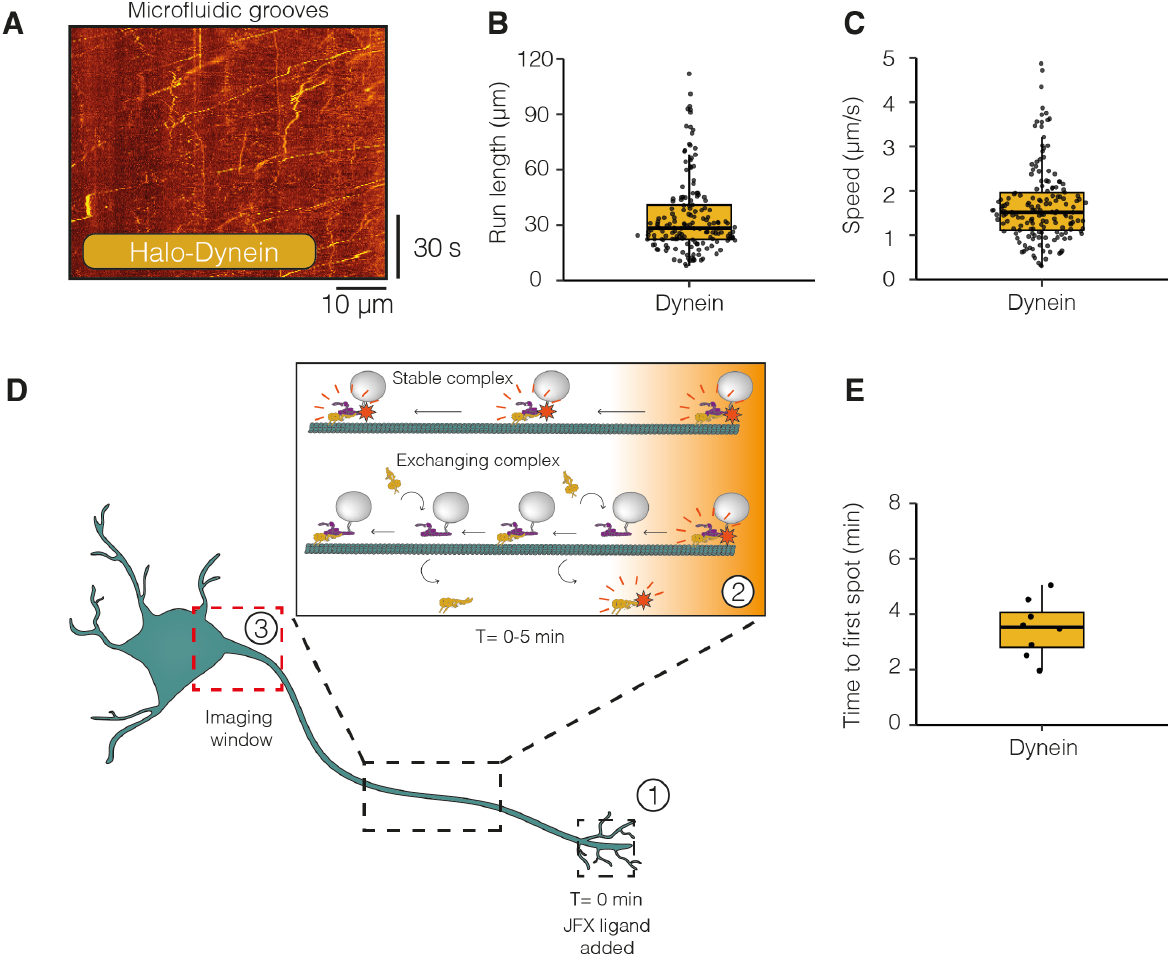
Dynein moves long-range along the axon in a stable complex. **A)** Example kymograph of retrograde dynein (Halo-*DYNC1H1*) movement in 21-23 DPI neurons treated with either JFX 554 or JFX 650. **B)** The run lengths of retrograde dynein particles in 21-23 DPI neurons. **C)** The speed of retrograde dynein particles in 21-23 DPI neurons (162 tracks, 21 videos, N=6). **D)** Schematic of experimental setup to explore dynein movement in the axon. 1: JFX dyes were added to the axon tip at T = 0 mins. This labels both diffusive and motile dynein. 2: Motile dynein moves along the axon. Dynein either forms a stable interaction with cargo and moves the entire length of the axon or undergoes short range movements and then dissociated from the complex. 3: Our imaging window at the proximal end of the axon. If dynein is stably bound, we expected to see the first fluorescent spot within 5 min. On the other hand, if dynein is exchanging on cargo, it should take longer. **E)** The amount of time until the first processive fluorescent particle was detected in dynein (Halo-*DYNC1H1*) 21-23 DPI neurons (8 videos, N=8). Boxplots shows median, first and third quartiles. Upper/lower whiskers extend to 1.5x the interquartile range.

To assess the level of dynein present in these moving spots we measured the distribution in intensities in a single 30 ms frame of the collected movies and compared them with the intensities of the static NEM-treated spots, described above, under the same imaging condition (Fig S3B). The distribution of intensities of static spots was between 0 and 200 AU above background, consistent with our photobleaching data suggesting 1-7 fluorophores with an average intensity of ∼26 AU (Fig 2F). The moving dyneins showed a narrower distribution between 0 and 100 AU consistent with between 1-4 fluorophores present per spot.

We observed run lengths of individual dynein spots of up to 110 µm, which is approximately the width of the imaging window. The average run length was shorter at 35.19 ± 0.66 µm, although this appears to be limited by the labelled spots going in and out of focus. The use of an endogenous HaloTag on dynein thus allows us to image much longer runs than observed previously^4,6^. However, with this setup we were unable to conclusively determine if the dyneins we observed had travelled the whole length of the axon.

To address this, we repeated the experiment but started imaging immediately after treatment with the halo ligand. If a dynein molecule is stably attached to the cargo, given the average speed observed (∼1.76 µm/s, Fig 3B), we would expect the first fluorescent dynein molecule to take under 5 min to traverse the 500 µm microfluidic groove. Alternatively, if dynein was rapidly exchanging on cargo and only moving short distances, we would expect a much slower arrival time of the first signal (Fig 3D). We saw the first fluorescent dynein come through within minutes (3.49 ± 0.12 min, Fig 3E). This suggests that at least some dynein is capable of stably binding cargos and moving them in a highly processive manner along the whole length of the axon.

### LIS1 and NDEL1 undergo long-range retrograde movements along the axon

We next asked if other dynein associated components also undergo long-range movement. Dynactin is required for dynein to bind cargo-specific, coiled-coil adaptors and to move processively^3^. We would therefore expect it to also travel long distances given our observations with dynein. However, the expectations are less clear for LIS1 and NDEL1. These proteins are both key in the formation of a motile dynein complex^3^, yet whether they are present on moving cargo in cells is unknown. *In vitro* studies with Lis1 have come to opposing conclusions about whether it can comigrate with dynein/dynactin complexes^23–27^. To directly visualise these associated proteins, we used our dynactin cell line (Halo-*ACTR10*, hiPSC) and generated cell lines with tagged LIS1 (*PAFAB1H1*-Halo, hESCs) and Ndel1 (Halo-*NDEL1*, hESCs). We then analysed retrograde movement of these proteins in 21-23 DPI iNeurons.

We saw multiple highly processive retrograde events not only with dynactin but also with LIS1 and NDEL1 (Fig 4A, Video 8-10). Although it appeared that there were fewer processive events for both LIS1 and NDEL1 than with dynein and dynactin, the difference was only statistically significant for NDEL1 (LIS1: 1.16 ± 0.12 min^-1^, NDEL1: 0.57 ± 0.11 min^-1^ vs dynein: 3.00 ± 0.27 min^-1^ and dynactin: 3.29 ± 0.66 min^-1^, Fig 4B). To understand if these components had also travelled the length of the axon, we repeated our experiment by measuring how long it took to visualise the first retrograde fluorescent particle to travel through the microfluidic grooves. We found that dynactin, LIS1 and NDEL1 all travelled through the groove in a similar time frame to dynein (Fig S4A). This suggests that like dynein these proteins bind to a cargo stably throughout their transport along an axon.

**Figure 4:**
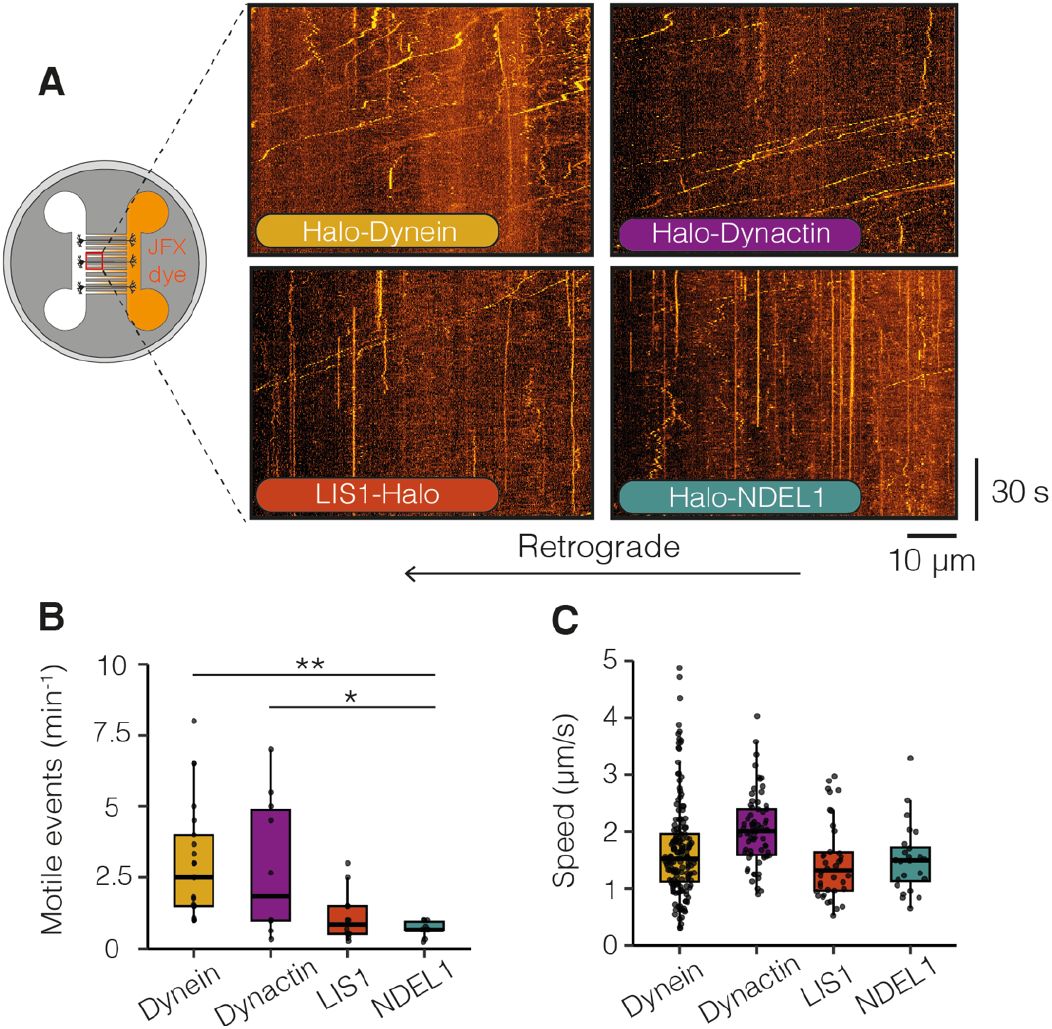
Dynein machinery moves long range retrogradely along the axon. **A)** Schematic of microfluidic device showing treatment in the axonal compartment with JFX halo ligand and imaging in the somatodendritic compartment. Example kymographs of retrograde dynein (Halo-*DYNC1H1*), dynactin (Halo-*ACTR10*), LIS1 (*PAFAH1B1*-Halo) and NDEL1 (Halo-*NDEL1*) movement in 21-23 DPI neurons. **B)** The frequency of retrograde motile events in dynein, dynactin, LIS1 and NDEL1 in 21-23 DPI neurons (dynein: 162 tracks, 21 videos, N=6; dynactin: 68 tracks, 10 videos, N=4; LIS1: 38 tracks, 14 videos, N=6; NDEL1: 24 tracks, 10 videos, N=3). Dynein vs NDEL1: **p= 0.0088, dynactin vs NDEL1: *p=0.0263, Kruskal-Wallis test, Dunn post hoc test. **C)** Graph showing the speed of retrograde dynein, dynactin, LIS1 and NDEL1 particles in 21-23 DPI neurons. Boxplots shows median, first and third quartiles. Upper/lower whiskers extend to 1.5x the interquartile range.

Previously, some *in vitro* studies showed that when LIS1 is present on dynein complexes it reduces their velocity compared to those without LIS1^23^. If this is the case in the axon, we would expect a LIS1 spot to have a lower average velocity compared to dynein in our iNeurons. However, we saw no significant difference in speed or pausing kinetics between any of the dynein machinery (Fig 4D, S4B, S4C). This suggests either that LIS1 is unable to have the same effect on the dynein motor in the cell as was seen *in vitro*, or alternatively that LIS1 is travelling on cargos without directly interacting with the motors driving transport.

### Dynein and dynactin reach the distal tip of the axon at different speeds

Many organelles are known to move bidirectionally^1^ and co-purify with both kinesin and dynein motors^28–31^. Therefore, we expected that both dynein and dynactin should be present on multiple kinesin-driven anterograde vesicles. To directly test this we treated dynein, dynactin, LIS1 and NDEL1 iNeurons with 200 nM JFX 554/650 in the somatodendritic compartment and imaged them in the axonal compartment. We observed anterograde movements for all components of the dynein machinery analysed (Fig 5A, Video 11-14), although there were significantly fewer LIS1 particles compared to dynein and dynactin (LIS1: 1.05 ± 0.10 min^-1^, dynein: 2.43 ± 0.15 min^-1^, dynactin: 2.87 ± 0.22 min^-1^ and NDEL1: 1.76 ± 0.21 min^-1^, Fig 5B).

**Figure 5:**
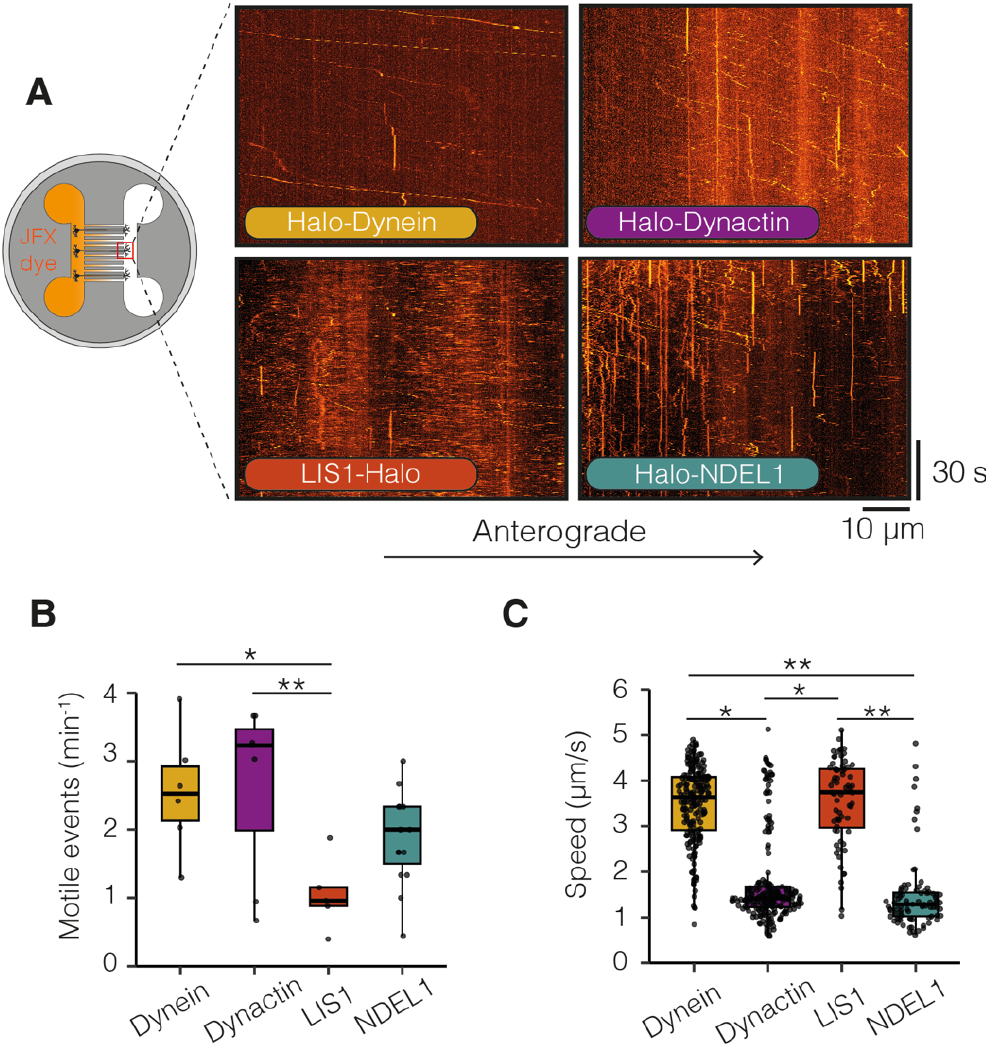
Dynein and its machinery travel to the distal tip of the axon separately. **A)** Schematic of microfluidic device showing treatment in the somatodendritic compartment with JFX 554/650 ligand and imaging in the axonal compartment. Example kymographs of retrograde dynein (Halo-*DYNC1H1*), dynactin (Halo-*ACTR10*), LIS1 (*PAFAH1B1*-Halo) and NDEL1 (Halo-*NDEL1*) movement in 21-23 DPI neurons. **B)** The frequency of anterograde motile events in dynein, dynactin, LIS1 and NDEL1 in 21-23 DPI neurons (dynein: 198 tracks, 6 videos, N=5; dynactin: 211 tracks, 7 videos, N=5; LIS1: 68 tracks, 5 videos, N=5; NDEL1: 86 tracks, 15 videos, N=3). Dynein vs LIS1: *p= 0.033, dynactin vs LIS1: **p=0.0077, Kruskal-Wallis test, Dunn post hoc test. **C)** The speed of anterograde dynein, dynactin, LIS1 and NDEL1 particles in 21-23 DPI neurons (dynein: 198 tracks, 6 videos, N=5; dynactin: 211 tracks, 7 videos, N=5; LIS1: 68 tracks, 5 videos, N=5; NDEL1: 86 tracks, 15 videos, N=3). Dynein vs dynactin: *p=0.044; dynein vs NDEL1: **p=0.0056; dynactin vs LIS1: *p=0.015; LIS1 vs NDEL1: **p=0.0018. Kruskal-Wallis test, Dunn post hoc test. Boxplot shows median, first and third quartiles. Upper/lower whiskers extend to 1.5x the interquartile range.

Strikingly, we found that the majority of dynein and LIS1 particles travelled at a similar speed that was significantly faster than that of dynactin and NDEL1 (dynein: 3.47 ± 0.04 µm/s and LIS1: 3.67 ± 0.07 µm/s vs dynactin: 1.78 ± 0.03 µm/s and NDEL1: 1.47 ± 0.01 µm/s, Fig 5C). We also observed that LIS1 and dynein show significantly fewer pauses during their anterograde transport than dynactin and NDEL1 (dynein: 5.25 ± 0.36% and LIS1: 3.30 ± 0.28% vs dynactin: 15.04 ± 0.33% and NDEL1: 20.13 ± 2.04%, Fig S5A, B). Given the difference in speed and pausing kinetics, it appears dynein and LIS1 are being transported to the axon tip via a different mechanism to dynactin and NDEL1.

Our dynactin line was made in a different stem cell background (hiPSCs) from the dynein, NDEL1 and LIS1 (hESCs). Although iNeurons differentiated from these two stem cells showed no significant difference in anterograde and retrograde transport velocities of membrane cargo and CTB (Fig S5C, D), we wanted to ensure the cell background was not the cause in the speed difference between dynein and dynactin. We therefore used a dual line with both dynein (Halo-*DYNC1H1*) and dynactin (*DCTN4*-SNAP) tagged in the same hESC line. We treated these iNeurons with either 200 nM JFX 554 or 1 µM SNAP-SiR to label dynein or dynactin respectively in the somatodendritic compartment. We again found that the majority of dynein moved significantly faster than dynactin in the anterograde direction (dynein: 3.61 ± 0.36 µm/s, dynactin: 1.88 ± 0.28 µm/s, Fig S5E). Overall, our data suggests that many dynein and LIS1 molecules are being trafficked to the distal tip separately to dynactin and NDEL1.

## Discussion

### Dynein moves long-range in the axon

Our data suggest that dynein is capable of long-range movement. Previous work using mouse hippocampal neurons reported runs of GFP-labelled dynein of up to 15 µm^4^. Here, due to the fact we can label dyneins at the axon tip and image them as they reach the cell body, we show they can move the entire length of the axon. In contrast to our work, a recent study suggested dynein moves cargo by multiple short runs, detaching after each movement^6^. However, transport in HeLa cells differs from neurons^16^, with membrane vesicles undergoing multiple rounds of short movements interspersed with pauses.

Determining how many dyneins are present on a certain cargo has proven challenging in cells. Previous work *in vitro* has suggested teams of dynein are required to move organelles^32,33^. In this study, we estimated that moving dynein spots that had traversed the axon contained 1-4 fluorophores. If fully labelled with the HaloTag this would correspond to 1-2 dynein molecules. In agreement with this, a recent study found 1-2 dyneins present on endosomes^6^. Other work has suggested that ∼8 dyneins might be required to move lysosomes in the cell ^34^. One potential explanation for the difference in numbers might be the size of cargo. The nature of our experiments selects for the fastest cargos in the axon and we know from previous work that smaller cargos, such as endosomes, which have room for fewer motors, move faster^16^ (Fig 1D). Another explanation could be that some of dynein dissociates from our cargos during transit. Despite this, it is clear at least some dyneins remain on cargos throughout transport along the axon.

Some vesicles are known to mature and change their composition during their transport in the axon. Examples are endosomes and autophagosomes, which slowly become more acidic as they turn into degradative organelles^35,36^. Previous work on autophagosomes suggested that different cargo-specific adaptors are required for dynein-driven transport in different segments of the axon^37^. An interesting question is how these observations relate to very long-distance movement of dynein. One possibility is that additional dynein molecules are tethered onto cargos independently of the activating adaptor. In this way the pool of dynein that moves along the axon would engage different adaptors when required. Another possibility is that some cargos engage dynein for the whole duration of their transport whereas others, which we would not detect in the experiments reported here, show exchange of both motors and adaptors.

### LIS1 and NDEL1 move retrogradely

LIS1 and NDEL1 are integral to dynein mediated trafficking^38^. Yet, it is only recently that their roles in this process have begun to be understood. LIS1 helps initiate dynein movement^23–25^ by disrupting its autoinhibition and supporting the formation of active complexes with dynactin^23,25,39–41^. On the other hand, NDEL1 recruits LIS1 to dynein ^42,43^. Whether these proteins remain part of the motile complex has been unclear. Some data suggest LIS1 co-migrates with dynein/dynactin complexes *in vitro*^26,27,44^, whereas others studies find LIS1 dissociates from moving complexes^23,25,41,45^. However, this issue had not been addressed in mammalian cells. Our data reveal that both LIS1 and NDEL1 are transported long distances along the axon. This raises the question of why they co-migrate with cargos. One possibility is that vesicles contain both actively engaged dynein/dynactin and a pool of reserve motors. In this case the presence of LIS1 would allow formation of new active dynein/dynactin complexes during a cargo’s journey along the axon. New initiation events may be required when cargos encounter obstacles. This was highlighted by a previous study where LIS1 and NDEL1 were shown to facilitate increased force production of dynein when cargo movement was restrained by an optical trap^46^. Overall, our study has revealed that both proteins are transported along the length of the axon and likely play a key role in long-range trafficking.

### Dynein and its machinery move to the axon tip at different speeds

The speed profiles of dynein, dynactin, LIS1 and NDEL1 moving in the anterograde direction fit into two groups: one containing dynein and LIS1, which travels much faster (∼3.5 µm/s) than the other containing dynactin and NDEL1 (∼1.6 µm/s). This raises the question of how these groups are transported at different speeds. Dynein, dynactin and LIS1 have all been found to bind the growing end of microtubules^47,48^. However, the speed of neuronal microtubule polymerization is ∼0.1 µm/s^16^ and therefore cannot account for any of the speeds we observed. Instead, it is likely the two groups are both driven by kinesin motors.

Kinesin-1, -2 and -3 are the major anterograde motors present in neurons and are responsible for trafficking the majority of organelles, proteins and RNA^2^. Previous work reported that certain kinesins are faster than others, with kinesin-3 being roughly 3 times faster than kinesin-1^49^. Therefore, dynein/LIS1 and dynactin/NDEL1 may be driven by different kinesins. Another mechanism would be to alter the speed of the same kinesin when it is bound to different cargos. For example, neuronal APP (amyloid precursor protein) vesicles, which are driven by kinesin-1, move much faster (∼3.6 µm/s) than other kinesin-1 cargos^50^. This faster speed depends on the presence of the adaptor protein JIP1^51^ and matches that of our anterograde moving dynein and LIS1.

### Implications of transporting dynein and dynactin separately to the axon tip

Several lines of evidence suggest kinesin and dynein are both present on cargos and can alternate activity to cause rapid reversals in direction^52^. These include colocalization of both motors on cargo^28,29^, the ability of both motors to simultaneously bind to the same adaptor proteins^30,31,53^ and observations that inhibition of either motor leads to bi-directional transport defects^29,54–56^. As dynactin is known to be required for dynein function, we had assumed that it travels with dynein on cargos in the anterograde direction. Although we saw a small percentage dynactin moving anterogradely at the same speed as dynein, the majority moved slower. This implies that many anterograde cargos lack at least one of the two main components of the dynein/dynactin machinery. What could explain this separation of dynein and dynactin?

One possibility is that the missing component, either dynein or dynactin, is picked up in transit and directly results in a reversal. Other examples of reversals due to motor recruitment are known. For example, in *U. maydis* kinesin-driven cargos reverse direction when they run into a dynein cargo moving in the other direction. In this case however the reversal is likely due to recruitment of both dynein and dynactin^57^. More recently, work in HeLa cells suggested dynactin, adaptors and cargos wait on microtubules and only move when dynein is recruited^6^, although the situation may be different from that in neurons where cargos are moving much longer distances. A second possibility is that dynein can allow a cargo to reverse without dynactin being present. Although it is widely accepted that dynactin is required for dynein function^3^ there have been reports of dynactin independent functions ^58,59^. This, however, does not explain the anterograde cargos that only have dynactin on them.

An alternative explanation is that neurons separate these proteins for rapidly delivery of the retrograde transport machinery to the axon tip. Previous work suggested ∼90% of dynein molecules are transported by a slow axonal transport driven by transient direct interactions between dynein and kinesin-1, which wafts it toward the axon tip^4^. However, calculations show that this slow delivery of dynein contributes relatively little to the population of retrogradely moving motors^60^. Our observations of a large flux of anterograde dynein and dynactin movement are consistent with these calculations. The separate movement of dynein and dynactin would have the advantage that they are less likely to be active and interfere with kinesin-driven transport. Likewise, the separate movement of the initiation factors NDEL1 and LIS1 would ensure that the retrograde transport machinery predominantly assembles where it is needed at the axon tip. Taken together, this ensures that neurons have an efficient transport system with microtubule motors only active when needed.

## Data availability

All data and analysis files are available at 10.5281/zenodo.8082407. The scripts used for photobleaching analysis can be found at https://github.com/carterlablmb.

## Acknowledgements

We thank J. O’Neil for suggesting iNeuron as model system; University of Cambridge, Mark Kotter lab for the provision of the neuron inducible human pluripotent stem cell line (NGN2-OPTi-OX); S. Bullock for helpful discussions, reading and help conceiving the project. R. Wademan and G. Manigrasso for help with cell culture; S.Chaaban for critical reading of the manuscript; J. Grimmett and T. Darling for providing scientific computing resources; and finally we thank the light microscope core facility at the MRC laboratory of molecular biology for technical assistance. This work was supported by Wellcome (210711/Z/18/Z), the Medical Research Council, as part of UK Research and Innovation (MRC file reference number MC_UP_A025_1011). For the purpose of open access, the author has applied a CC BY public copyright license to any author-accepted manuscript version arising.

## Author contributions

A.D.F performed the experiments and analysis and prepared the figures. A.D.F, M.B, T.B and A.B generated the knock-in cell lines used in this manuscript. A.D.F and A.P.C conceived the project and wrote the manuscript.

## Online Methods

### Human stem cell culture and NGN2 neuronal differentiation

hESC (H9 line; WiCELL) and hiPSC (Bit Bio Ltd)^7^ that harbour a doxycycline-inducible NGN2 transgene in the AAVS1 locus were kept on Cultrex basement membrane extract (35 µg/cm^2^, R&D systems) and fed every other day with mTeSR plus (STEMCELL Technologies). Cells were kept at 37°C in a 5% CO2 incubator.

In order to differentiate into iNeurons, cells were dissociated into single cells with accutase (STEMCELL Technologies) and 300,000 were cells plated per well of Cultrex coated 6-well dish. For the first 24 hr, cells were kept in mTeSR plus and CloneR (1X, STEMCELL Technologies). After that, media was switched to differentiation media (DMEM/F12 (Gibco), GlutaMAX (1X, Gibco), Non-Essential Amino Acids (1X, Gibco), N2 supplement (1X, Gibco), Penicillin/Streptomycin (1%) and doxycycline (1 µg/ml, Merck)) for 48 hr. After this time, cells were dissociated with accutase and immediately plated into microfluidics (PDMS mould on glass bottom dish (HBST-5040, #1.5H, 0.005 mm, Willco Wells)) containing neuronal media (Neurobasal (Gibco), GlutaMAX (1X), B27 supplement (Gibco), BDNF (10 ng/ml, Peprotech), NT3 (10 ng/ml, Peprotech), Penicillin/Streptomycin (1%) and doxycycline (1 µg/ml)). Microfluidics were coated with poly-D-lysine (20 µg/ml, Merck) and Geltrex hESC-Qualified reduced growth factor basement membrane matrix (0.12-0.18 mg/ml, ThermoFisher). A 25% media exchange took place every two days until 7 DPI where doxycycline was removed from the neuronal media. iNeurons were cultured until 21-23 DPI when imaging took place. Cells were kept at 37°C in a 5% CO2 incubator.

### CRISPR knock-in of HaloTag to stem cells

Knock-in of the HaloTag to hESC or hiPSCs was done following established methods^61^. Briefly, ribonucleoprotein (RNP) complexes were formed with 1 µl HiFi Cas9 (4 µg/µl, IDT), 6 µl synthetic sgRNA (30 µM, with the following protospacer sequences ***DYNC1H1***: CTCCGACATGGTGTCGCGCT, ***ACTR10***: CGTAGAGCGGCATGGTAGTA, ***PAFAB1H1***: GCCGTTGATTGTGTCTCCTT, ***NDEL1***: TTCACAGGCTTTCTTGATCA, ***DCTN4***: CCCTCCAGTGGAACCTT, Synthego) and nucleofection buffer P3 (Lonza). ssDNA (6 µg) containing 100-150 nt homology arms flanking the HaloTag of SNAP-tag coding sequence were added. 215,000 accutase-dissociated hESC or hiPSC were mixed with RNP complexes and nucleofected with the 4D-Nucleofector (Lonza, CA-137). The cells were then plated in a 6-well dish coated with rhLaminin-521 (0.5 µg/cm^2^, Gibco) with mTeSR plus and CloneR. Cells were left until confluent (∼7 days). At this time point, cells were treated with 200 nM JF646 HaloTag ligand (Promega) for 20 min. Cells were then accutase-dissociated and washed twice in 4 ml PBS. Halo-positive cells were then flow-sorted and 3000 Halo+ cells were plated at clonal density on a Cultrex-coated 10 cm dish. Cells were kept in mTeSR plus and CloneR. Individual colonies were picked and screened for successful gene editing by PCR and sanger sequencing. Knock-in lines were then differentiated into iNeurons following the above protocol.

### Immunofluorescence

iNeurons at 21-23 DPI were fixed in 4% paraformaldehyde (Sigma) in PBS for 12 min at room temperature. Cells were then washed with PBS, permeabilised and blocked for 15 min in permeabilisation buffer (0.5% bovine serum albumin, 10% donkey serum, 0.2% Triton X-100 in PBS). Primary antibodies against SMI-31P (BioLegend, 801601, 1:500), MAP2 (synaptic systems, 188004, 1:500) and *β*3-tubulin (Sigma, T2200, 1:1000) were diluted in blocking buffer (0.5% BSA, 10% DS in PBS) and incubated with cells for 1 hr at room temperature. iNeurons were then washed three times with PBS and incubated with the correct fluorescently conjugated secondary antibody (Invitrogen, 1:1000) diluted in blocking buffer for 1 hr. Cells were washed with PBS and mounted with ProLong Diamond antifade mountant (ThermoFisher). Finally, iNeurons were imaged using an inverted Zeiss LSM 780 using a 63x, 1.4 NA DIC Plan-Apochromat oil immersion objective.

### Photobleaching step analysis

Halo-*DYNC1H1* and Halo-*ACTR10* iNeurons were cultured in microfluidics until 21-23 DPI. Cells were treated with 1 nM JFX Halo ligand in the axonal chamber for 20 mins. Ligand was washed out with new neuronal medium and cells were left overnight at 37°C in a 5% CO2 incubator. Cells where then treated with 0.5 µM N-ethyl maleimide (NEM, ThermoFisher) for 20 min. Cells were imaged using an inverted Nikon 100x, 1.49 NA CFI Apochromat oil immersion TIRF lens. In order to improve signal to noise, cells were imaged using HILO imaging^6^. HILO settings were optimised for each condition by altering the angle of incidence of the excitation laser (561 nm or 640 nm) between 57°-60°. Laser power was kept constant at 50% (15 mW at fiber, LU-N4, Nikon). Time-lapse images were acquired at 30 Hz with 30 ms exposure (sCMOS, 95% QE, Prime 95b, Teledyne Photometrics) continuously for 2 min. Spots were picked using ImageJ and intensity analysis was run using custom scripts in Matlab at https://github.com/carterlablmb.

### Live cell imaging

Live imaging of iNeurons took place between 21-23 DPI. Endosome, lysosome and mitochondrial transport was assessed in microfluidics with the addition of either 1 µg/ml CTB AlexaFluor 488, 50nM Lysotracker Deep Red or 100 nM Mitotracker Deep Red FM to the axonal compartment, or Vybrant DiD membrane dye (2000x, all ThermoFisher) in the somatodendritic compartment for 30 min at 37°C. Cells were washed and then new pre-warmed low fluorescent BrainPhys (STEMCELL Technologies) supplemented with GlutaMAX (1X, Gibco), B27 supplement (1X, Gibco), BDNF (10 ng/ml, Peprotech), NT3 (10 ng/ml, Peprotech), Penicillin/Streptomycin (1%) was added to cells. 15 min later, transport was imaged at 37°C using an inverted Zeiss LSM 880 using a 63x, 1.4 NA DIC Plan-Apochromat oil-immersion objective. Images were taken at 2 Hz over a period of 2-4 min. For figure S5, CTB and Vybrant DiD membrane dye were imaged by using an inverted Nikon 100x, 1.49 NA CFI Apochromat oil immersion TIRF lens. In order to improve signal to noise, cells were imaged using HILO imaging^6^. HILO settings were optimised for each condition by altering the angle of incidence of the excitation laser (561 nm or 640 nm) between 57°-60°. Laser power was kept constant at 50% (15 mW at fiber, LU-N4, Nikon). Time lapse images were acquired at 2 Hz with 30 ms exposure (sCMOS, 95% QE, Prime 95b, Teledyne Photometrics) for 3 min.

For endogenous HaloTag/Snap-tag imaging, iNeuron media was exchanged to pre-warmed low fluorescent BrainPhys medium (STEMCELL Technologies) supplemented with GlutaMAX (1X, Gibco), B27 supplement (1X, Gibco), BDNF (10 ng/ml, Peprotech), NT3 (10 ng/ml, Peprotech), Penicillin/Streptomycin (1%). We first treated cells with increasing concentrations of Halo ligand (1-500 nM) to assess which led to the best labelling. We determined that 200 nM was sufficient. Therefore, cells were treated with either 200 nM JFX 554/650 ^62^ in the axonal (Retrograde: Fig 3, S3, 4, S4), or 200 nM JFX 554/650 or 1 µM SNAP-SiR in the somatodendritic compartment (Anterograde: Fig 5, S5). Cells were imaged immediately at either the somatodendritic compartment (Retrograde: Fig 3, S3, 4, S4) or axonal compartment (Anterograde: Fig 5, S5) of the microfluidic at 37°C (stage-top incubator, Oko Labs). This was done using an inverted Nikon 100x, 1.49 NA CFI Apochromat oil immersion TIRF lens. In order to improve signal to noise, cells were imaged using HILO imaging^6^. HILO settings were optimised for each condition by altering the angle of incidence of the excitation laser (561 nm or 640 nm) between 57°-60°. Laser power was kept constant at 50% (15 mW at fiber, LU-N4, Nikon). Time lapse images were acquired at 2 Hz with 30 ms exposure (sCMOS, 95% QE, Prime 95b, Teledyne Photometrics) for 2-15 min.

### Analysis and quantification

Image analysis was done using Fiji (NIH,MD). In order to quantify organelle and endogenous Halo-tagged protein kinetics, Trackmate imaging software was used ^63,64^. For this analysis, spots were tracked using the manual tracking implementation and output data consisted of spot, edge and track files. The edges files contain data of each individual movement whereas the track files summarise the overall movement of the particles. Only spots that moved over 10 µm were tracked. Pauses were defined as the particles moving slower than 0.1 µm/s.

### Statistical analysis

All analysis was undertaken in R^65^. Data was assessed for normality by Shapiro-Wilk test. To assess the difference between two groups with a normal distribution a student’s T-test was used. For analysis of multiple groups in which the data was not normally distributed, the Kruskal-Wallis test was used followed by the Dunn test for multiple comparisons. All statistics were done on the mean value from each biological replicate. Statistical significance is noted as follows: * p ≤ 0.05, ** p ≤ 0.01, and *** p ≤ 0.001. All statistical tests and associated p values are indicated in figure legend.

## Supplemental figures

**Figure 1 supplemental:**
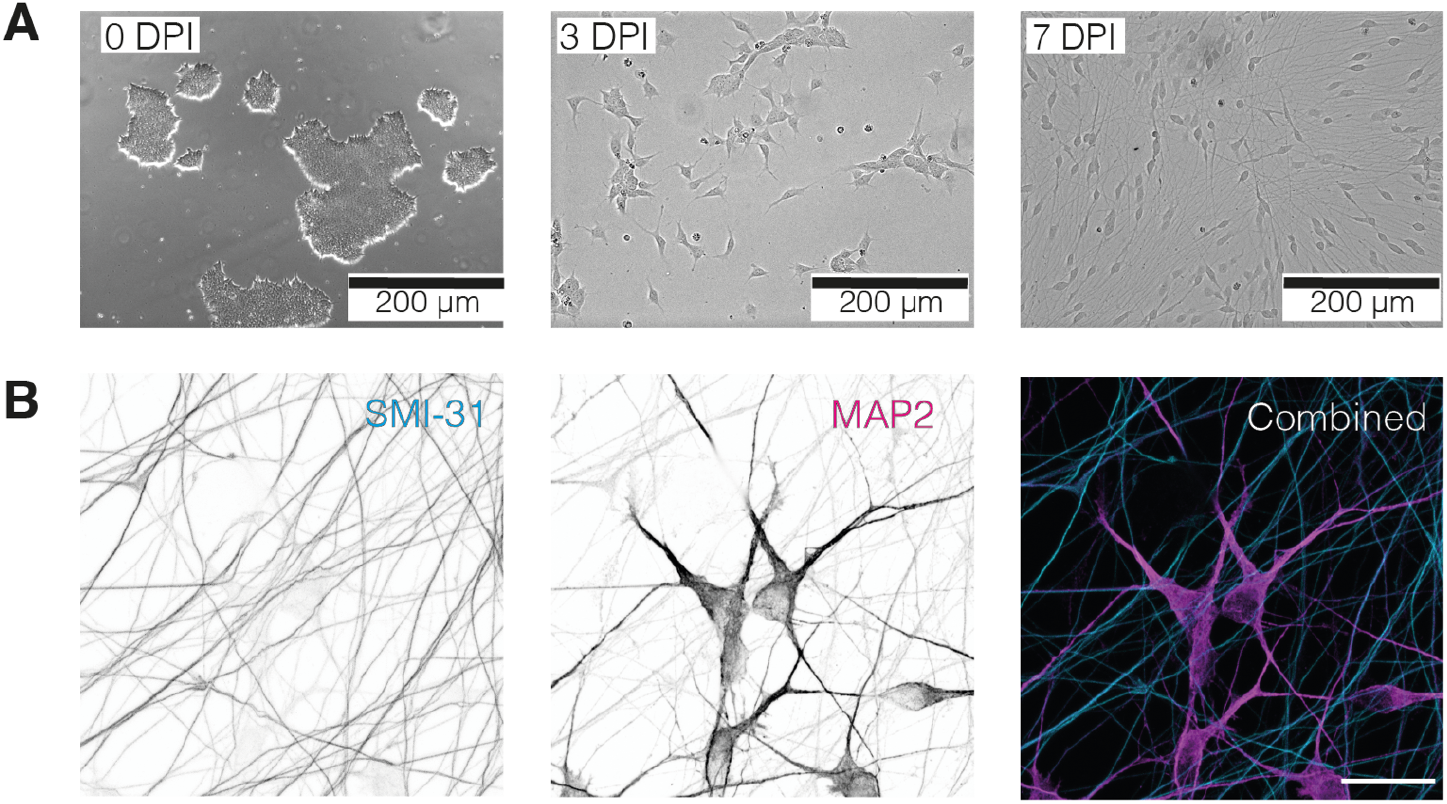
iNeurons as a model to study dynein mediated transport. **A)** Example immunofluorescence images of different stages of iNeuron differentiation. DPI 0 = stems cells, DPI 3 = 3 days post induction with doxycycline, DPI 7 = 7 days post induction with doxycycline. **B)** Image of 21-23 DPI iNeurons showing staining with axonal (SMI-31, cyan) and dendritic (MPA2, magenta) markers. Scale bar is 20 µM.

**Figure 2 supplemental:**
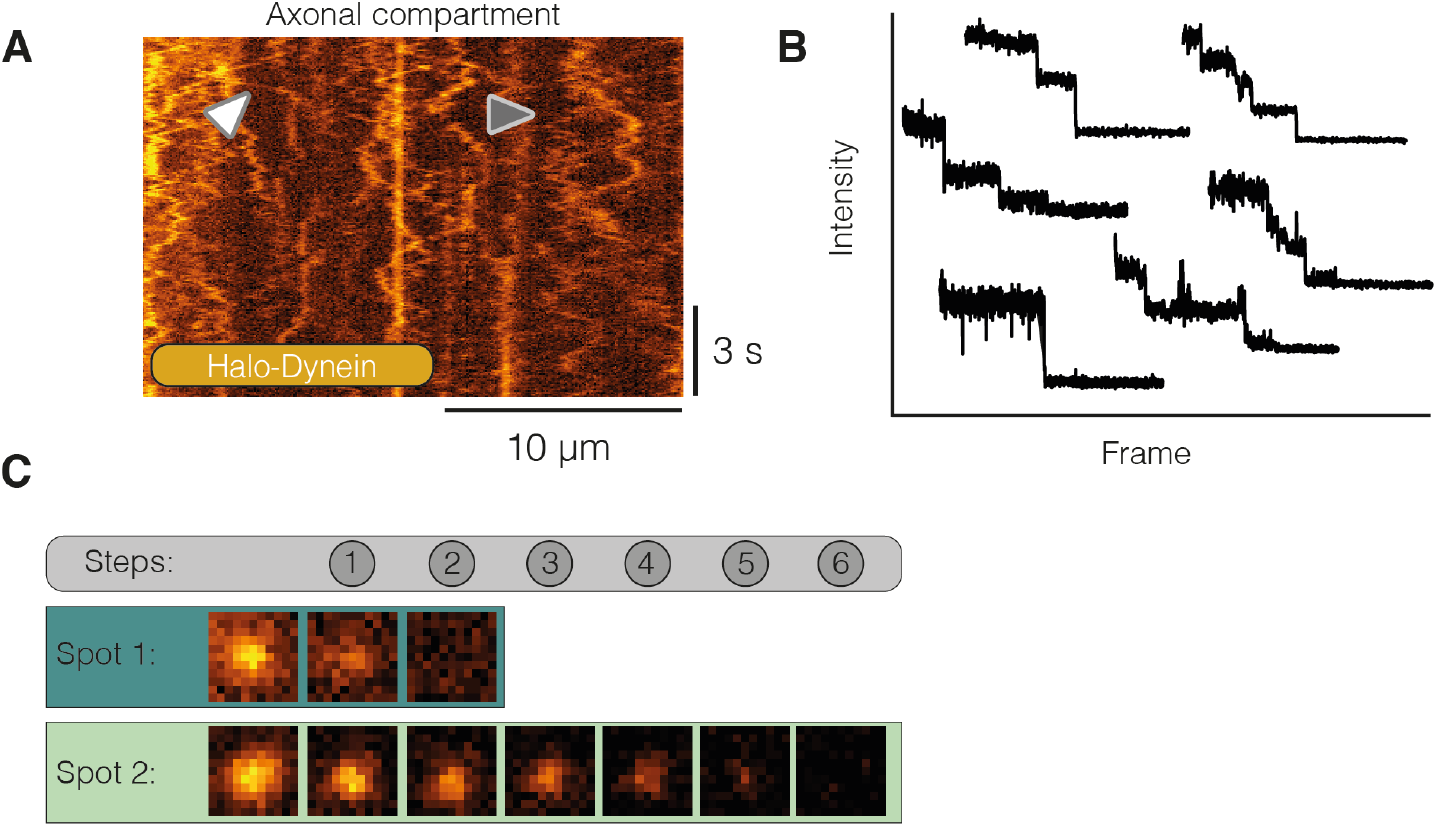
Visualising dynein in iNeurons. **A)** Example kymograph of 21-23 DPI Halo-dynein treated with 1 nM JFX 554. White arrow highlights processive event. Grey arrow shows diffusive motility. **B)** 6 bleaching traces from dynein spots. The traces were separated to enhance clarity. **C)** Images of two spots during bleaching from Fig 2B, C. The teal spot undergoes 2 bleaching steps and corresponds to the bleaching trace in Fig 2B. The light green spot undergoes 6 bleaching steps and corresponds to Fig 2C.

**Figure 3 supplemental:**
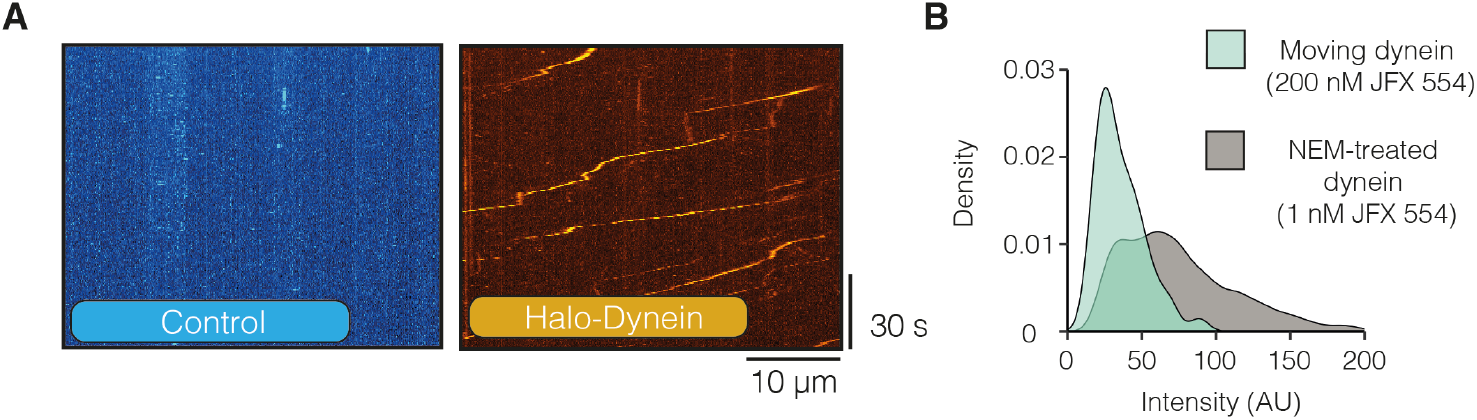
Dynein moves long-range along the axon in a stable complex. **A)** Example kymographs of control (untagged) and Halo-*DYNC1H1* 21-23 DPI iNeurons treated with 500 nM JFX 554. **B)** The intensity of moving dynein spots vs dynein spots treated with NEM (Fig 2). Given the step size calculated in Fig 2F, the intensities correspond to between 1-4 fluorescent molecules.

**Figure 4 supplemental:**
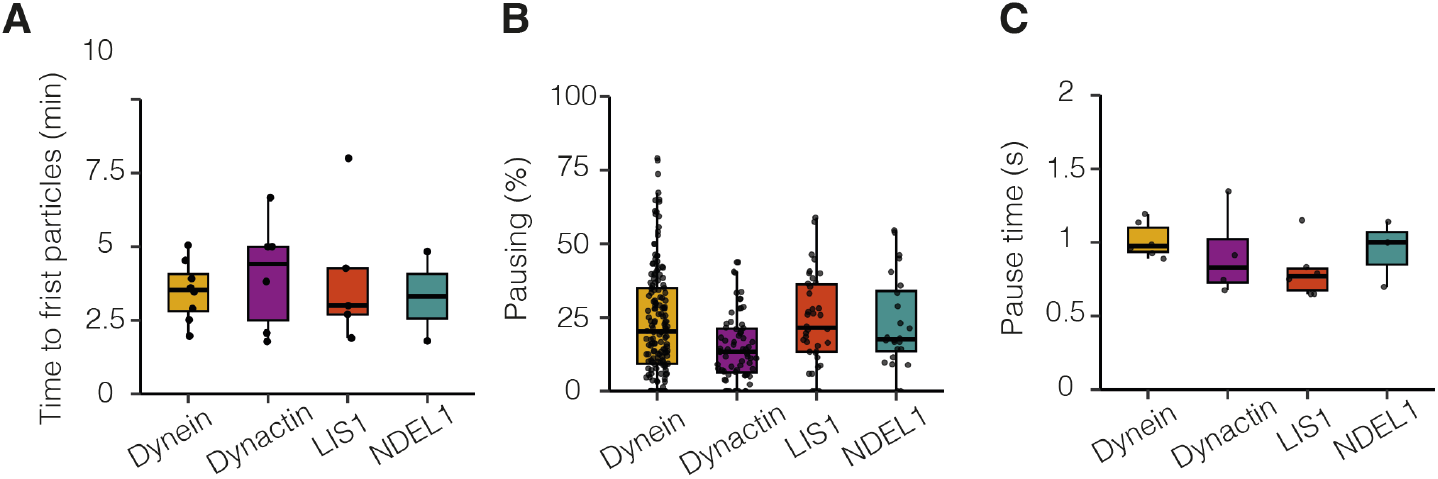
Dynein machinery moves long range retrogradely along the axon. **A)** The amount of time until the first processive fluorescent particle was detected in dynein, dynactin, LIS1 and NDEL1 21-23 DPI neurons (dynein: 8 videos, N=8; dynactin: 6 videos, N=6; LIS1: 5 videos, N=5; NDEL1: 2 videos, N=2). **B)** The amount of time dynein, dynactin, LIS1 and NDEL1 spent pausing in the retrograde direction (dynein: 162 tracks, 21 videos, N=6; dynactin: 68 tracks, 10 videos, N=4; LIS1: 38 tracks, 14 videos, N=6; NDEL1: 24 tracks, 10 videos, N=3). Pauses are defined as spots moving slower than 0.1 µm/s. **C)** The average length of time each pause lasted for dynein, dynactin, LIS1 and NDEL1. Boxplot shows median, first and third quartiles. Upper/lower whiskers extend to 1.5x the interquartile range.

**Figure 5 supplemental:**
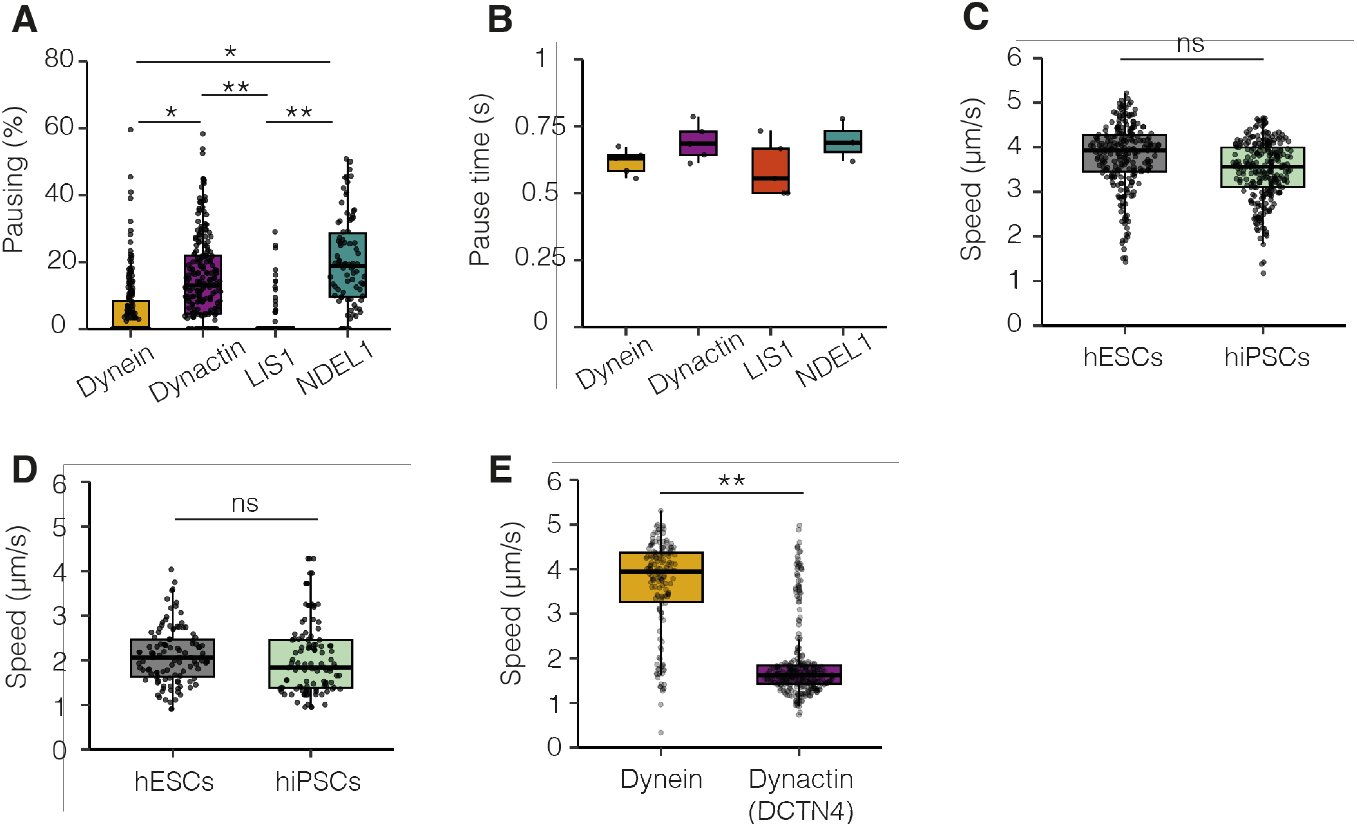
Dynein and its machinery travel to the distal tip of the axon separately. **A)** The percent of time dynein, dynactin, LIS1 and NDEL1 spent pausing. Dynein vs dynactin: *p=0.044; dynein vs NDEL1: *p=0.026; dynactin vs LIS1: **p=0.0037; LIS1 vs NDEL1: **p=0.0028. Kruskal-Wallis test, Dunn post hoc test. Pauses are defined as spots moving slower than 0.1 µm/s. **B)** Average length of time each pause lasted for dynein, dynactin, LIS1 and NDEL1 in the anterograde direction. **C)** The speed of anterograde cargos in 21-23 DPI iNeurons generated from either hESCs or hiPSCs (hESCs: 236 tracks, 3 videos, N=3; hiPSCs: 250 tracks, 3 videos, N=3). **D)** The speed of retrograde cargos in 21-23 DPI iNeurons generated from either hESCs or hiPSCs (hESCs: 111 tracks, 3 videos, N=3; hiPSCs: 107 tracks, 3 videos, N=3). **E)** The speed of anterograde dynein (Halo-*DYNC1H1*) and dynactin (*DCTN4*-SNAP) particles (dynein: 150 tracks, 9 videos, N=3; dynactin: 323 tracks, 12 videos, N=3). Dynein vs dynactin: **p=0.0047, student’s t-test. Boxplots shows median, first and third quartiles. Upper/lower whiskers extend to 1.5x the interquartile range.

## Supplemental table 1

**Table S1:**

| Test | diff | p adj |
| --- | --- | --- |
| Lyso:Anterograde-CTB:Anterograde | -0.03444457 | 0.9998031 |
| Mito:Anterograde-CTB:Anterograde | -0.92503693 | 0.0001869 |
| CTB:Retrograde-CTB:Anterograde | -<br>0.17573796 | 0.7747167 |
| Lyso:Retrograde-CTB:Anterograde | -0.67574519 | 0.0030455 |
| Mito:Retrograde-CTB:Anterograde | -0.86830289 | 0.0003401 |
| Mito:Anterograde-Lyso:Anterograde | -0.89059236 | 0.0002682 |
| CTB:Retrograde-Lyso:Anterograde | -0.14129339 | 0.8904110 |
| Lyso:Retrograde-Lyso:Anterograde | -0.64130062 | 0.0046199 |
| Mito:Retrograde-Lyso:Anterograde | -0.83385832 | 0.0004944 |
| CTB:Retrograde-Mito:Anterograde | 0.74929897 | 0.0012809 |
| Lyso:Retrograde-Mito:Anterograde | 0.24929174 | 0.4683607 |
| Mito:Retrograde-Mito:Anterograde | 0.05673404 | 0.9978115 |
| Lyso:Retrograde-CTB:Retrograde | -0.50000723 | 0.0268761 |
| Mito:Retrograde-CTB:Retrograde | -0.69256493 | 0.0024910 |
| Mito:Retrograde-Lyso:Retrograde | -0.19255770 | 0.7071265 |

## Videos

**Video 1:** Endosome transport (CTB) in DPI 21-23 iNeurons. Imaged at 1 fps. Playback 10 fps.

**Video 2:** Lysosome transport (Lysotracker) in DPI 21-23 DPI iNeurons. Imaged at 1 fps. Playback 10 fps.

**Video 3:** Mitochondrial transport (Mitotracker) in DPI 21-23 iNeurons. Imaged at 1 fps. Playback 10 fps.

**Video 4:** Imaging of Halo-*DYNC1H1* DPI 21-23 iNeurons treated with 1 nM JFX 650 in the axonal compartment. Imaged at 30 fps. Playback 60 fps.

**Video 5:** Photobleaching in Halo-*DYNC1H1* DPI 21-23 iNeurons in the axonal compartment treated with 1 nM JFX 554 and 0.5 µM NEM. Imaged at 30 fps. Playback 60 fps.

**Video 6:** Photobleaching in Halo-*ACTR10* DPI 21-23 iNeurons in the axonal compartment treated with 1 nM JFX 554 and 0.5 µM NEM. Imaged at 30 fps. Playback 60 fps.

**Video 7:** Imaging of Halo-*DYNC1H1* retrograde movement in DPI 21-23 iNeurons treated with 200 nM JFX 554. Imaged at 2 fps. Playback 20 fps.

**Video 8:** Imaging of Halo-*ACTR10* retrograde movement in DPI 21-23 iNeurons treated with 200 nM JFX 554. Imaged at 2 fps. Playback 20 fps.

**Video 9:** Imaging of Halo-*PAFAH1B1* retrograde movement in DPI 21-23 iNeurons treated with 200 nM JFX 554. Imaged at 2 fps. Playback 20 fps.

**Video 10:** Imaging of Halo-*NDEL1* retrograde movement in DPI 21-23 iNeurons treated with 200 nM JFX 554. Imaged at 2 fps. Playback 20 fps.

**Video 11:** Imaging of Halo-*DYNC1H1* anterograde movement in DPI 21-23 iNeurons treated with 200 nM JFX 554. Imaged at 2 fps. Playback 20 fps.

**Video 12:** Imaging of Halo-*ACTR10* anterograde movement in DPI 21-23 iNeurons treated with 200 nM JFX 554. Imaged at 2 fps. Playback 20 fps.

**Video 13:** Imaging of Halo-*PAFAH1B1* anterograde movement in DPI 21-23 iNeurons treated with 200 nM JFX 554. Imaged at 2 fps. Playback 20 fps.

**Video 14:** Imaging of Halo-*NDEL1* anterograde movement in DPI 21-23 iNeurons treated with 200 nM JFX 554. Imaged at 2 fps. Playback 20 fps.

## References

1. Maday, S., Twelvetrees, A. E., Moughamian, A. J. & Holzbaur, E. L. F. Axonal Transport: Cargo-Specific Mechanisms of Motility and Regulation. Neuron 84, 292–309 (2014).

2. Sleigh, J. N., Rossor, A. M., Fellows, A. D., Tosolini, A. P. & Schiavo, G. Axonal transport and neurological disease. Nat. Rev. Neurol. (2019). doi:10.1038/s41582-019-0257-2

3. Reck-Peterson, S. L., Redwine, W. B., Vale, R. D. & Carter, A. P. The cytoplasmic dynein transport machinery and its many cargoes. Nat. Rev. Mol. Cell Biol. 19, 382–398 (2018).

4. Twelvetrees, A. E. E. et al. The Dynamic Localization of Cytoplasmic Dynein in Neurons Is Driven by Kinesin-1. Neuron 90, 1000–1015 (2016).

5. Ha, J. et al. A neuron-specific cytoplasmic dynein isoform preferentially transports TrkB signaling endosomes. J. Cell Biol. 181, 1027–1039 (2008).

6. Tirumala, N. A. et al. Single-molecule imaging of cytoplasmic dynein in cellulo reveals the mechanism of motor activation and cargo capture. bioRxiv (2022). doi:10.1016/j.bpj.2021.11.2703

7. Pawlowski, M. et al. Inducible and Deterministic Forward Programming of Human Pluripotent Stem Cells into Neurons, Skeletal Myocytes, and Oligodendrocytes. Stem Cell Reports 8, 803–812 (2017).

8. Watson, E. T., Pauers, M. M., Seibert, M. J., Vevea, J. D. & Chapman, E. R. Synaptic vesicle proteins are selectively delivered to axons in mammalian neurons. Elife 12, 1–26 (2023).

9. Ananthanarayanan, V. et al. Dynein motion switches from diffusive to directed upon cortical anchoring. Cell 153, 1526 (2013).

10. Hulme, A. J., Maksour, S., St-Clair Glover, M., Miellet, S. & Dottori, M. Making neurons, made easy: The use of Neurogenin-2 in neuronal differentiation. Stem Cell Reports 17, 14–34 (2022).

11. Zhang, Y. et al. Rapid single-step induction of functional neurons from human pluripotent stem cells. Neuron 78, 785–798 (2013).

12. Schörnig, M. et al. Comparison of induced neurons reveals slower structural and functional maturation in humans than in apes. Elife 10, 1–78 (2021).

13. Boecker, C. A., Olenick, M. A., Gallagher, E. R., Ward, M. E. & Holzbaur, E. L. F. ToolBox: Live Imaging of intracellular organelle transport in induced pluripotent stem cell-derived neurons. Traffic 21, 138–155 (2020).

14. Wang, C. et al. Scalable Production of iPSC-Derived Human Neurons to Identify Tau-Lowering Compounds by High-Content Screening. Stem Cell Reports 9, 1221–1233 (2017).

15. Park, J. W., Vahidi, B., Taylor, A. M., Rhee, S. W. & Jeon, N. L. Microfluidic culture platform for neuroscience research. Nat. Protoc. 1, 2128–2136 (2006).

16. Fellows, A. D., Rhymes, E. R., Gibbs, K. L., Greensmith, L. &Schiavo, G. IGF 1R regulates retrograde axonal transport of signalling endosomes in motor neurons. EMBO Rep. 21, 1–16 (2020).

17. Kulkarni, V. V., Stempel, M. H., Anand, A., Sidibe, D. K. & Maday, S. Retrograde Axonal Autophagy and Endocytic Pathways Are Parallel and Separate in Neurons. J. Neurosci. 42, 8524–8541 (2022).

18. Los, G. V et al. HaloTag: A Novel Protein Labeling Technology for Cell Imaging and Protein Analysis. ACS Chem. Biol. 3, 373–382 (2008).

19. Banaz, N., Mäkelä, J. & Uphoff, S. Choosing the right label for single-molecule tracking in live bacteria: Side-by-side comparison of photoactivatable fluorescent protein and Halo tag dyes. J. Phys. D. Appl. Phys. 52, (2019).

20. Broadbent, D. G., Barnaba, C., Perez, G. I. & Schmidt, J. C. Quantitative analysis of autophagy reveals the role of ATG9 and ATG2 in autophagosome formation. J. Cell Biol. 222, (2023).

21. Pfister, K. K., Wagner, M. C., Bloom, G. S. & Brady, S. T. Modification of the Microtubule-Binding and ATPase Activities of Kinesin by N-Ethylmaleimide (NEM) Suggests a Role for Sulfhydryls in Fast Axonal Transport. Biochemistry 28, 9006–9012 (1989).

22. Scott, D. A., Das, U., Tang, Y. & Roy, S. Mechanistic Logic Underlying the Axonal Transport of Cytosolic Proteins. Neuron 70, 441–454 (2011).

23. Htet, Z. M. et al. LIS1 promotes the formation of activated cytoplasmic dynein-1 complexes. Nat. Cell Biol. 22, 518–525 (2020).

24. Qiu, R., Zhang, J. & Xiang, X. LIS1 regulates cargo-adapter-mediated activation of dynein by overcoming its autoinhibition in vivo. J. Cell Biol. 218, 3630–3646 (2019).

25. Elshenawy, M. M. et al. Lis1 activates dynein motility by modulating its pairing with dynactin. Nat. Cell Biol. 22, 570–578 (2020).

26. Baumbach, J. et al. Lissencephaly-1 is a context-dependent regulator of the human dynein complex. Elife 6, 1–31 (2017).

27. Gutierrez, P. A., Ackermann, B. E., Vershinin, M. & McKenney, R. J. Differential effects of the dynein-regulatory factor Lissencephaly-1 on processive dynein-dynactin motility. J. Biol. Chem. 292, 12245–12255 (2017).

28. Maday, S., Wallace, K. E. & Holzbaur, E. L. F. Autophagosomes initiate distally and mature during transport toward the cell soma in primary neurons. J. Cell Biol. 196, 407–417 (2012).

29. Encalada, S. E., Szpankowski, L., Xia, C. H. & Goldstein, L. S. B. Stable kinesin and dynein assemblies drive the axonal transport of mammalian prion protein vesicles. Cell 144, 551–565 (2011).

30. Fenton, A. R., Jongens, T. A. & Holzbaur, E. L. F. Mitochondrial adaptor TRAK2 activates and functionally links opposing kinesin and dynein motors. Nat. Commun. 1–15 (2021). doi:10.1038/s41467-021-24862-7

31. Canty, J., Hensley, A. M., Aslan, M., Jack, A. & Yildiz, A. Trak adaptors regulate the recruitment and activation of dynein and kinesin in mitochondrial transport. Nat. Commun. 122, 25a–26a (2023).

32. Rai, A. et al. Dynein Clusters into Lipid Microdomains on Phagosomes to Drive Rapid Transport toward Lysosomes. Cell 164, 722–734 (2016).

33. Rai, A. K., Rai, A., Ramaiya, A. J., Jha, R. & Mallik, R. Molecular adaptations allow dynein to generate large collective forces inside cells. Cell 152, 172–182 (2013).

34. Ori-Mckenney, K. M., Xu, J., Gross, S. P. & Vallee, R. B. A cytoplasmic dynein tail mutation impairs motor processivity. Nat. Cell Biol. 12, 1228–1234 (2010).

35. Kulkarni, V. V. & Maday, S. Neuronal endosomes to lysosomes : A journey to the soma. J. Cell Biol. 217, 2977–2979 (2018).

36. Cason, S. E. & Holzbaur, E. L. F. Selective motor activation in organelle transport along axons. Nature Reviews Molecular Cell Biology 23, 699–714 (2022).

37. Cason, S. E. et al. Sequential dynein effectors regulate axonal autophagosome motility in a maturation-dependent pathway. J. Cell Biol. 220, (2021).

38. Lam, C., Vergnolle, M. A. S., Thorpe, L., Woodman, P. G. & Allan, V. J. Functional interplay between LIS1, NDE1 and NDEL1 in dynein-dependent organelle positioning. J. Cell Sci. 123, 202–212 (2010).

39. Marzo, M. G., Griswold, J. M. & Markus, S. M. Pac1/LIS1 promotes an uninhibited conformation of dynein that coordinates its localization and activity. Nat. Cell Biol. (2020). doi:10.1101/684290

40. Qiu, R., Zhang, J. & Xiang, X. LIS1 regulates cargo-adapter-mediated activation of dynein by overcoming its autoinhibition in vivo. J. Cell Biol. 218, 3630–3646 (2019).

41. Gillies, J. P. et al. Structural Basis for Cytoplasmic Dynein-1 Regulation by Lis1. Elife 11, 1–29 (2022).

42. Okada, K. et al. Conserved Roles for the Dynein Intermediate Chain and Ndel1 in Assembly and Activation of Dynein. bioRxiv 1–22 (2023).

43. Garrott, S. R. et al. Ndel1 modulates dynein activation in two distinct ways. bioRxiv 1–23 (2023).

44. Jha, R., Roostalu, J., Cade, N. I., Trokter, M. & Surrey, T. Combinatorial regulation of the balance between dynein microtubule end accumulation and initiation of directed motility. EMBO J. 36, 3387–3404 (2017).

45. Egan, M. J., Tan, K. & Reck-Peterson, S. L. Lis1 is an initiation factor for dynein-driven organelle transport. J. Cell Biol. 197, 971–982 (2012).

46. Reddy, B. J. N. et al. Load-induced enhancement of Dynein force production by LIS1-NudE in vivo and in vitro. Nat. Commun. 7, (2016).

47. Carvalho, P., Gupta, M. L., Hoyt, M. A. & Pellman, D. Cell cycle control of kinesinmediated transport of Bik1 (CLIP-170) regulates microtubule stability and dynein activation. Dev. Cell 6, 815–829 (2004).

48. Vaughan, K. T., Tynan, S. H., Faulkner, N. E., Echeverri, C. J. & Vallee, R. B. Colocalization of cytoplasmic dynein with dynactin and CLIP-170 at microtubule distal ends. J. Cell Sci. 112, 1437–1447 (1999).

49. Lipka, J., Kapitein, L. C., Jaworski, J. & Hoogenraad, C. C. Microtubule-binding protein doublecortin-like kinase 1 (DCLK1) guides kinesin-3-mediated cargo transport to dendrites. Embo J. 1, 302–318 (2016).

50. Araki, Y. et al. The novel cargo Alcadein induces vesicle association of kinesin-1 motor components and activates axonal transport. EMBO J. 26, 1475–1486 (2007).

51. Tsukamoto, M. et al. The cytoplasmic region of the amyloid β-protein precursor (APP) is necessary and sufficient for the enhanced fast velocity of APP transport by kinesin-1. FEBS Lett. 592, 2716–2724 (2018).

52. Hancock, W. O. Bidirectional cargo transport: Moving beyond tug of war. Nat. Rev. Mol. Cell Biol. 15, 615–628 (2014).

53. Kendrick, A. A. et al. Hook3 is a scaffold for the opposite-polarity microtubule-based motors cytoplasmic dynein-1 and KIF1C. J. Cell Biol. 218, 2982–3001 (2019).

54. Martin, M. et al. Cytoplasmic Dynein, the Dynactin Complex, andKinesin Are Interdependent and Essential for Fast Axonal Transport. Mol. Biol. Cell 10, 3717–3728 (1999).

55. Sainath, R. & Gallo, G. The dynein inhibitor Ciliobrevin D inhibits the bidirectional transport of organelles along sensory axons and impairs NGF-mediated regulation of growth cones and axon branches. Dev. Neurobiol. 75, 757–777 (2015).

56. Ally, S., Larson, A. G., Barlan, K., Rice, S. E. & Gelfand, V. I. Opposite-polarity motors activate one another to trigger cargo transport in live cells. 187, 1071–1082 (2009).

57. Bielska, E. et al. Hook is an adapter that coordinates kinesin-3 and dynein cargo attachment on early endosomes. J. Cell Biol. 204, 989–1007 (2014).

58. McKenney, R. J., Weil, S. J., Scherer, J. & Vallee, R. B. Mutually exclusive cytoplasmic dynein regulation by NudE-Lis1 and dynactin. J. Biol. Chem. 286, 39615–39622 (2011).

59. Wadsworth, P. & Lee, W. L. Microtubule motors: Doin’ it without dynactin. Curr. Biol. 23, R563–R565 (2013).

60. Susalka, S. J., Hancock, W. O. & Pfister, K. K. Distinct cytoplasmic dynein complexes are transported by different mechanisms in axons. Biochim. Biophys. Acta - Mol. Cell Res. 1496, 76–88 (2000).

61. Bruntraeger, M., Byrne, M., Long, K. & Bassett, R. Editing the Genome of Human Induced Pluripotent Stem Cells Using CRISPR/Cas9 Ribonucleoprotein Complexes. Methods in Molecular Biology 1961, (2019).

62. Grimm, J. B. et al. A General Method to Improve Fluorophores Using Deuterated Auxochromes. JACS Au 1, 690–696 (2021).

63. Tinevez, J. Y. et al. TrackMate: An open and extensible platform for single-particle tracking. Methods 115, 80–90 (2017).

64. Ershov, D. et al. TrackMate 7: integrating state-of-the-art segmentation algorithms into tracking pipelines. Nat. Methods 19, 829–832 (2022).

65. R. R: A language and environment for statistical computing. R Foundation for Statistical Computing, Vienna, Austriae. (2014).

